# Interleg coordination is not strictly controlled during walking

**DOI:** 10.1101/2023.01.24.525466

**Authors:** Takahiro Arai, Kaiichiro Ota, Tetsuro Funato, Kazuo Tsuchiya, Toshio Aoyagi, Shinya Aoi

**Affiliations:** Graduate School of Informatics, Kyoto University, Yoshida-Honmachi, Sakyo-ku, Kyoto 606-8501, Japan; Cybozu, Inc., 2-7-1 Nihombashi, Chuo-ku, Tokyo 103-6027, Japan; Department of Mechanical Engineering and Intelligent Systems, Graduate School of Informatics and Engineering, The University of Electro-Communications, 1-5-1 Choufugaoka, Choufu, Tokyo 182-8585, Japan; Department of Aeronautics and Astronautics, Graduate School of Engineering, Kyoto University, Kyoto daigaku-Katsura, Nishikyo-ku, Kyoto 615-8540, Japan; Department of Mechanical Science and Bioengineering, Graduate School of Engineering Science, Osaka University, 1-3 Machikaneyama, Toyonaka, Osaka 560-8531, Japan

## Abstract

In human walking, the left and right legs move alternately, half a stride out of phase with each other. Although various parameters, such as stride frequency, stride length, and duty factor, vary with walking speed, the antiphase relationship of the leg motion remains unchanged. This is the case even during running. However, during walking in left-right asymmetric situations, such as walking with unilateral leg loading, walking along a curved path, and walking on a split-belt treadmill, the relative phase between left and right leg motion shifts from the antiphase condition to compensate for the asymmetry. In addition, the phase relationship fluctuates significantly during walking of elderly people and patients with neurological disabilities, such as those caused by stroke or Parkinson’s disease. These observations suggest that appropriate interleg coordination is important for adaptive walking and that interleg coordination is strictly controlled during walking of healthy young people. However, the control mechanism of interleg coordination remains unclear. In the present study, we derive a quantity that models the control of interleg coordination during walking of healthy young people by taking advantage of a state-of-the-art method that combines big data science with nonlinear dynamics. This is done by modeling this control as the interaction between two coupled oscillators through the phase reduction theory and Bayesian inference method. However, the results were not what we expected. Specifically, we found that the relative phase between the motion of the legs is not actively controlled until the deviation from the antiphase condition exceeds a certain threshold. In other words, the control of interleg coordination has a dead zone like that in the case of the steering wheel of an automobile. Such forgoing of control presumably enhances energy efficiency and maneuverability during walking. Furthermore, the forgoing of control in specific situations, where we expect strict control, also appears in quiet standing. This suggests that interleg coordination in walking and quiet standing have a common characteristic strategy. Our discovery of the dead zone in the control of interleg coordination not only provides useful insight for understanding gait control in humans, but also should lead to the elucidation of the mechanisms involved in gait adaptation and disorders through further investigation of the dead zone.

## Introduction

During human walking, the left and right legs move alternately, half a stride out of phase with each other [2]. In general, various locomotion parameters, such as gait frequency, stride length, and duty factor, change if the gait speed varies. However, the antiphase relationship of the leg motion remains unchanged. This is true even if the gait pattern changes to running. By contrast, during walking in which there is left-right asymmetry with regard to the body or environment, such as walking with unilateral leg loading [13], walking along a curved path [7], or walking on a split-belt treadmill [26], the relative phase between the legs shifts from the antiphase condition to compensate for the asymmetry. In addition, the phase relationship fluctuates significantly during walking of elderly people and patients with neurological disabilities, such as those caused by stroke or Parkinson’s disease [23, 24, 25, 29, 30]. These findings indicate that appropriate relative phase (that is, appropriate interleg coordination) is important for adaptive walking. This seems to suggest that the relative phase is strictly controlled during walking of healthy young people, with the relative phase quickly returning to the antiphase condition after being perturbed away from it.

To understand interleg coordination during human walking, it has been investigated how the relative phase between the motion of the left and right legs depends on the situations described above. However, it remains largely unclear to what extent it is strictly controlled in each situation. This is mainly because there are limitations on the degree to which the control of interleg coordination can be quantitatively elucidated due to the complexity of neural dynamics, musculo-skeletal dynamics, and interactions with the environment during walking. Elucidating the control of interleg coordination would not only clarify the adaptive strategy in human walking, but also contribute to the development of rehabilitation techniques for persons with gait disorders and gait assistive devices, such as robotic exoskeletons [1, 6, 12].

The present study aims to quantitatively clarify the control of interleg coordination during human walking by taking advantage of both data science and dynamical systems theory, employing the Bayesian inference method [4, 20] and phase reduction theory [9, 14, 17, 31]. Specifically, we model the rhythmic motion of the left and right legs by two coupled limit-cycle oscillators and describe these dynamics using two coupled phase oscillators. The phase coupling function between the two oscillators represents the nature of the control of the relative phase between the motion of the legs, i.e., interleg coordination. To identify this phase coupling function in human walking, we first measured the leg motion during walking of subjects on a treadmill with intermittent perturbations in the belt speed. Next, from the measured time-series data, we derived the phase coupling function using the Bayesian inference method. We analyze the characteristics of the control of interleg coordination during human walking on the basis of this function.

## Result

We measured kinematic data through observation of eight healthy men (Subjects A–H) during walking on a treadmill whose belt speed was intermittently disturbed from 1.0 m/s for a short time (Fig. 1(a)). More precisely, the belt speed was increased or decreased by 0.6 m/s over a period of 0.1 s, and then returned to 1.0 m/s over a period of 0.1 s. This resulted in the sudden collapse of the antiphase relationship of the leg motion and allowed us to investigate how the relative phase of the leg motion recovered after being disturbed. Each trial lasted approximately 60 s and contained ten disturbances during which the speed was varied. These disturbances were separated by intervals of approximately 5 s, which was sufficiently long to allow the return to normal steady-state walking. We regard the belt-speed disturbance, written *p*(*t*), as an external perturbation. We used three types of perturbation conditions for each trial: acceleration, deceleration, and mixed conditions. Only acceleration and deceleration perturbations of the belt speed were applied under the acceleration and deceleration conditions, respectively, while both perturbations were applied randomly under the mixed condition. We calculated the angle between the line connecting the greater trochanter of the hip and the head of the second metatarsal of the foot and a vertical line for each leg from the measured kinematic data. There angles are denoted by *φ*_L_(*t*) and *φ*_R_(*t*), where the subscripts L and R indicate the left leg and right leg, respectively.

**Figure 1:**
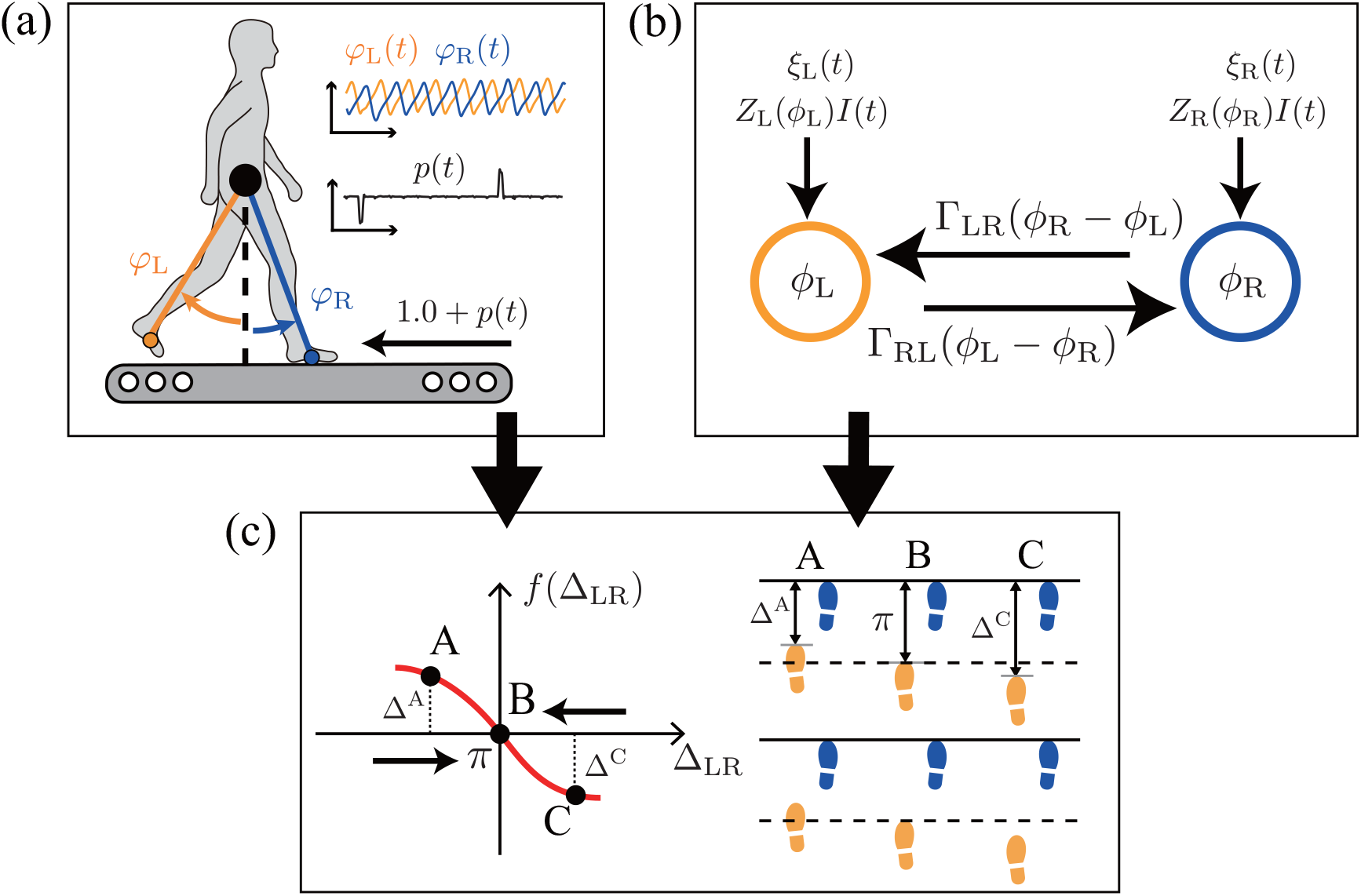
Schematic diagram of the study. **(a)** Measurement of the leg motion, represented by *φ_i_*(*t*) (*i* ∈ (L, R)), during walking on a treadmill with a perturbation in the belt speed, represented by *p*(*t*). **(b)** Two coupled limit-cycle oscillators with phases *ϕ_i_*(*t*) (*i* ∈ (L, R)) whose dynamics are described by phase equations with phase coupling functions Γ_*ij*_ (*ϕ_j_* – *ϕ_i_*), phase sensitivity functions *Z_i_*(*ϕ_i_*), external perturbation *I*(*t*), and Gaussian noise *ξ_i_* ((*i, j*) = (L, R), (R, L)). **(c)** Integration of the measured data and phase equations to derive the dynamics of the relative phase Δ_LR_, which reflects the control of interleg coordination. When Δ_LR_ = *π*, corresponding to B, the legs move in an exactly alternating manner. As Δ_LR_ moves away from *π*, the deviation from this antiphase relationship increases (see A and C, with Δ_LR_ = Δ^A^ and Δ^C^). If the interleg coordination is strictly controlled to maintain the antiphase relationship, the function *f*(Δ_LR_), which describes the control of interleg coordination, should intersect the horizontal axis at *π* with a steep negative slope.

We model the motion of the left and right legs by two coupled limit-cycle oscillators, whose phases are *ϕ*_L_ and *ϕ*_R_ ∈ (0, 2*π*], and describe these dynamics using phase equations on the basis of the phase reduction theory [9, 14, 17, 31] as follows (Fig. 1(b)):

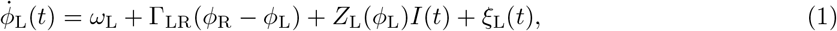

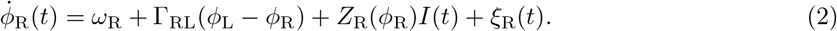

Here, *ω*_L_ and *ω*_R_ are the natural frequencies, Γ_LR_ and Γ_RL_ are phase coupling functions, *Z*_L_ and *Z*_R_ are phase sensitivity functions, *I* is an external perturbation (rescaled from *p*(*t*)), and *ξ*_L_ and *ξ*_R_ are independent Gaussian white noise terms that represent experimental uncertainty, which satisfy 〈(*ξ_i_*(*t*)〉 = 0 and 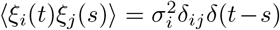 for (*i, j*) = (L, R) and (R, L), where *σ_i_* is the intensity of the Gaussian noise *ξ_i_*, and *δ_ij_* and *δ*(*t*) are the Kronecker and Dirac delta functions, respectively. We determined *ϕ_i_*(*t*) (*i* ∈ (L, R)) from the measured time series of the kinematic angle *φ_i_*(*t*), and also determined *I*(*t*) from the measured time series of the external perturbation *p*(*t*). We determined the best values for *ω_i_*, Γ_*ij*_, and *Z_i_* ((*i, j*) = (L, R), (R, L)) from the time series of *ϕ_i_*(*t*) and *I*(*t*) using the Bayesian inference method [17, 19, 20, 21]. We did the same for *σ_i_* in order to determine the extent to which Eqs. (1) and (2) are capable of modeling the dynamics of the observed human walking. We deem that a small intensity indicates that our deterministic model properly describes the dynamics and that the influence of the stochastic process is small. In previous studies [19, 20, 21], *Z_i_I* was not used in the identification of the phase equation, and the parameter values were obtained using data measured after external perturbations were applied. However, including *Z_i_I* in this identification allowed us to determine the parameter values more accurately through use of data measured while external perturbations were being applied.

Using phase equations derived through the procedure described above, we investigate how the relative phase between the motion of the legs, i.e., interleg coordination, is controlled during human walking. Specifically, if we ignore the phase response to the external perturbation in Eqs. (1) and (2) and focus on the transient dynamics after the perturbation is applied, we obtain the following equation for the relative phase Δ_LR_ ≔ *ϕ*_R_ – *ϕ*_L_:

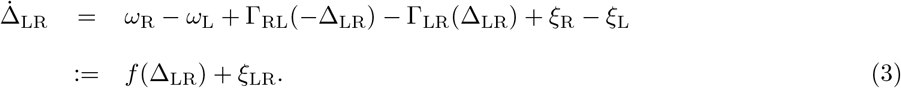

Here, we have *ξ*_LR_ ≔ *ξ*_R_ – *ξ*_L_, which satisfies 〈*ξ*_LR_(*t*)〉 = 0 and 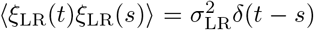 where 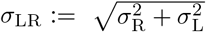, and *f*(Δ_LR_) describes the deterministic dynamics of Δ_LR_, i.e., the control of interleg coordination. If the antiphase relationship, Δ_LR_ = *π*, is tightly maintained, it is required that *f*(Δ_LR_) be 0 and have a steep negative slope at Δ_LR_ = *π* (Fig 1(c)).

Figure 2 displays the time series of *φ_i_*(*t*), *ϕ_i_*(*t*), Δ_LR_(*t*), and *p*(*t*) (*i* ∈ (L, R)) for one representative subject (Subject G). There, an increase of *ϕ_i_*(*t*) by 2*π* represents one cycle of *φ_i_*(*t*). It is seen that Δ_LR_(*t*) is greatly disturbed in response to the external perturbation (highlighted in green) and then returns to the neighborhood of *π*, the antiphase value. However, it does not converge to *π* but rather fluctuates about this value. This suggests that the dynamics of the relative phase possesses Lyapunov stability about *π*, not asymptotic stability. We now proceed to elucidate how *f*(Δ_LR_) reflects this stability.

**Figure 2:**
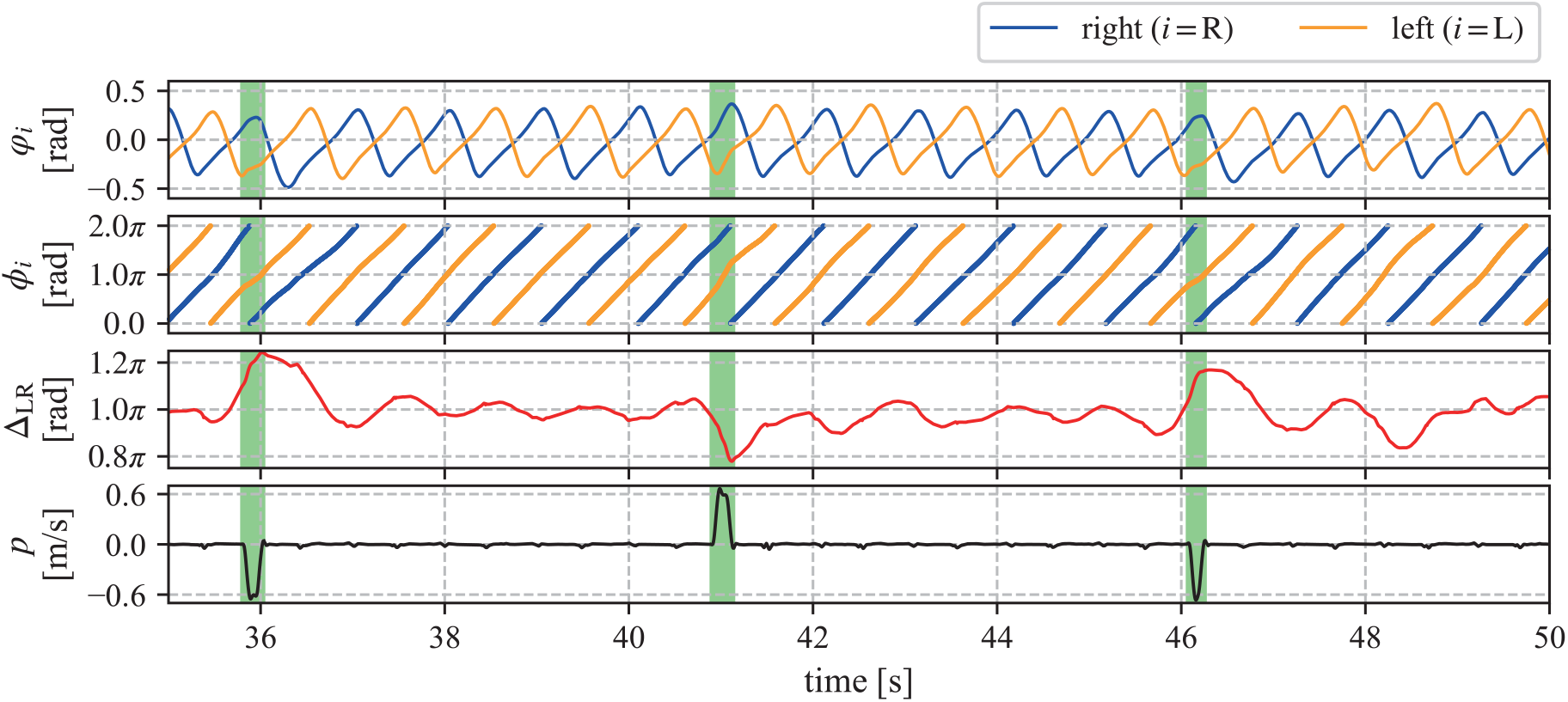
Measured and reconstructed time series for one representative subject (Subject G) under mixed conditions. The leg motion, represented by *φ_i_*(*t*) (*i* ∈ (L, R)), measured during walking was transformed to the oscillator phases *ϕ_i_*(*t*) (*i* ∈ (L, R)), from which we obtain the relative phase Δ_LR_. The green regions indicate the times at which the measured external perturbation, *p*(*t*), is large.

Figures 3(a) and 3(b) display the functions *Z_i_*(*i* ∈ (L, R)) and *f* (Δ_LR_), respectively, evaluated from the data for one representative subject (Subject G). It is seen that both *Z*_L_ and *Z*_R_ possess unimodal shapes with peaks near *π* and are close to zero in the range from 0 to *π*/2 for all perturbation conditions. Because Δ_LR_ fluctuates within a narrow region around *π*, as shown in Fig. 2, there was not a sufficient amount of data outside this region for the evaluation of *f*(Δ_LR_). Therefore, we limited our evaluation of *f*(Δ_LR_) to a range of 3 standard deviations around the mean of the observed Δ_LR_. Although we expected that *f*(Δ_LR_) would intersect the horizontal axis at one point near *π* with a steep negative slope, in accordance with the strict control of interleg coordination, as shown in Fig. 1(c), the results were not what we expected for any of the perturbation conditions. Specifically, we found that although *f*(Δ_LR_) has a steep negative slope sufficiently far from *π*, it is close to zero near *π*. Thus, we found that *f*(Δ_LR_) has a flat region with a value close to 0 in the neighborhood of the antiphase condition. This result indicates that the relative phase between the legs is not actively controlled until the deviation from the antiphase condition exceeds a certain threshold, where the antiphase relationship is lost as shown in Fig. 3(c). We obtained similar results from the other subjects (see Supplementary Information A). These characteristic properties of *f*(Δ_LR_) remained even when we excluded *Z_i_I* from the phase equation in the determination of *f*(Δ_LR_) (see Supplementary Information B). However, the form of *f*(Δ_LR_) obtained with *Z_i_I* exhibits these characteristics more clearly than that without *Z_i_I*.

**Figure 3:**
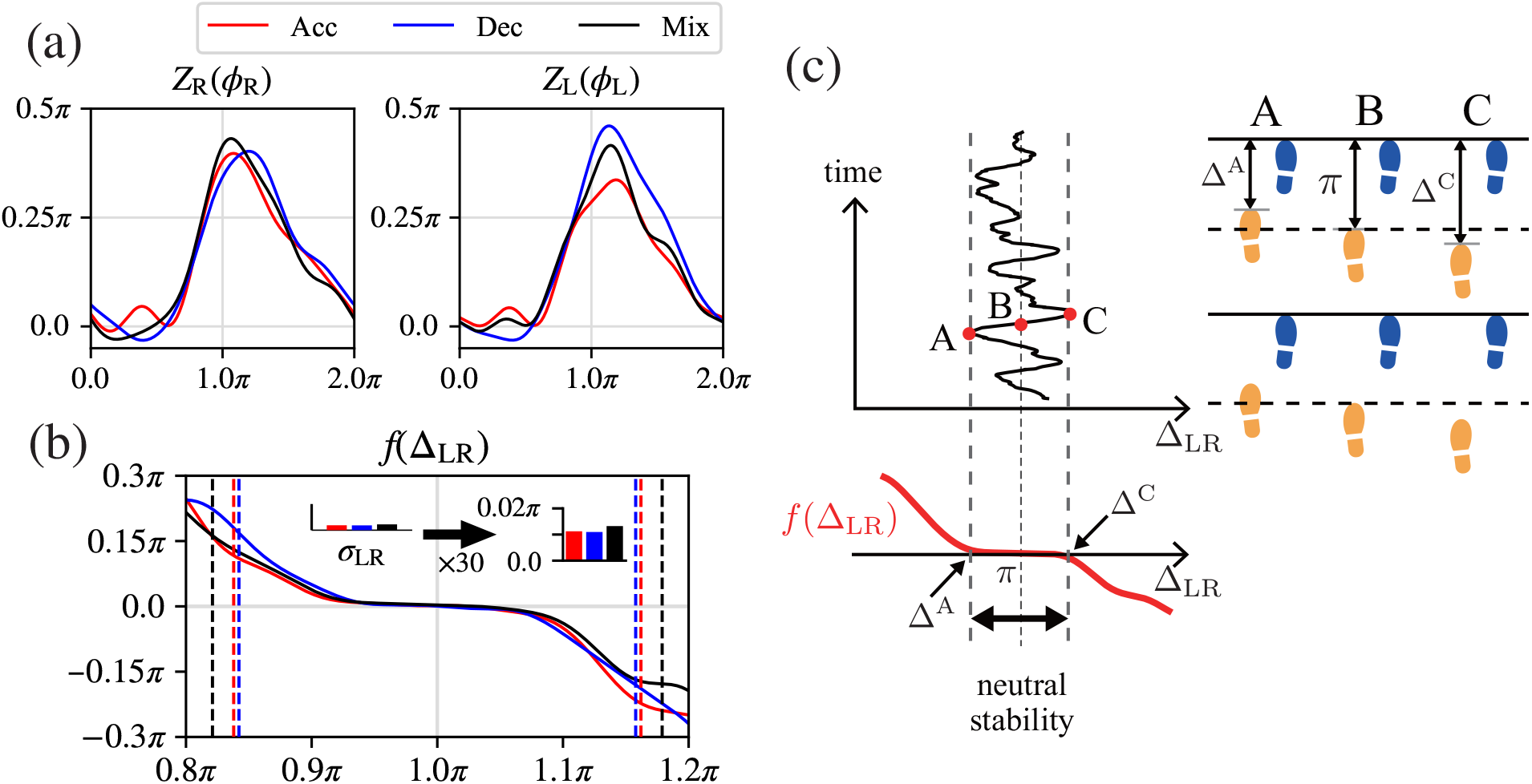
Results for the control of interleg coordination for one representative subject (Subject G). **(a)** Phase sensitivity functions *Z_i_*(*ϕ_i_*) (*i* ∈ (L, R)) for acceleration (Acc), deceleration (Dec), and mixed (Mix) conditions. **(b)** Control of interleg coordination, represented by *f*(Δ_LR_). The vertical dotted lines indicate 3 standard deviations from the mean of the observed Δ_LR_. The function *f*(Δ_LR_) is shifted in order to place the mean of the observed values of Δ_LR_ at *π* to improve visualization. The histogram displays the noise intensity, σ_LR_. (An expanded histogram, with values 30 times larger than the actual values, is also given.) The mean and standard deviation of the derived noise intensity for all results are 0.0106*π* and 0.0021*π*, respectively. These values are too small to cause deviation of Δ_LR_ from the flat region of *f*(Δ_LR_). **(c)** A schematic diagram for the interpretation of *f*(Δ_LR_). The flat region near Δ_LR_ = *π* at B (neutral stability) and the steep negative slope in the regions away from *π* reflect the fact that the relative phase between the legs is not actively controlled until the deviation from *π* exceeds a threshold (Δ^A^ – *π* at A and Δ^C^ – *π* at C), where motion of the legs deviates significantly from the antiphase relationship.

To clarify the universality of our findings in the control of interleg coordination represented by *f*(Δ_LR_), specifically, the existence of a flat region in which *f*(Δ_LR_) is close to 0 in the neighborhood of Δ_LR_ = *π*, and steep negative slopes outside this flat region (Fig. 3(c)), we compared the results for *f*(Δ_LR_) among the subjects. For this purpose, we approximated *f*(Δ_LR_) by a piecewise linear function to characterize these properties within 3 standard deviations around the mean of the observed Δ_LR_, as shown in Fig. 4(a). There, *l*_C_ is the length of the flat region, which reflects the extent to which the deviation from the antiphase condition is ignored, and *g*_L_ and *g*_R_ are the slopes in the regions to the left and right of the flat region, respectively, which reflect how rapidly the deviation outside the flat region attenuates. These three parameters, *l*_C_, *g*_L_, and *g*_R_, were determined to minimize the discrepancy between the original and approximated functions using the grid search method under the condition that the two functions coincide at the left and right endpoints of the approximated function. When there is no flat region in the original function, *l*_C_ = 0 is satisfied. Figure 4(b) displays the approximate forms of *f*(Δ_LR_) for the original forms appearing in Fig. 3(b). (Similar results for all subjects appear in Supplementary Information C.) Figure 4(c) presents the mean and standard deviation of *l*_C_ among the subjects. It is seen that *l*_C_ is approximately 0.4 and the deviation among the subjects is small for all perturbation conditions. We confirmed that l_C_ is significantly larger than 0 using a one-tailed *t*-test (*p* < 0.01). Indeed, the mean value of *l*_C_ is much larger than its standard deviation, and we thus conclude that *l*_C_ has a positive value. Figure 4(d) presents the mean and standard deviation of *g*_L_ and *g*_R_ among the subjects. It is seen that both |*g*_L_| and |*g*_R_| are approximately 2.0, sufficiently large that deviations rapidly attenuate for all perturbation conditions. We also confirmed that |*g*_L_| and |*g*_R_| are significantly larger than 0 using a one-tailed *t*-test (*p* < 0.01). These results confirm that *f*(Δ_LR_) has a flat region with a value close to 0 in the vicinity of Δ_LR_ = *π* and steep negative slopes outside this region for all perturbation conditions.

**Figure 4:**
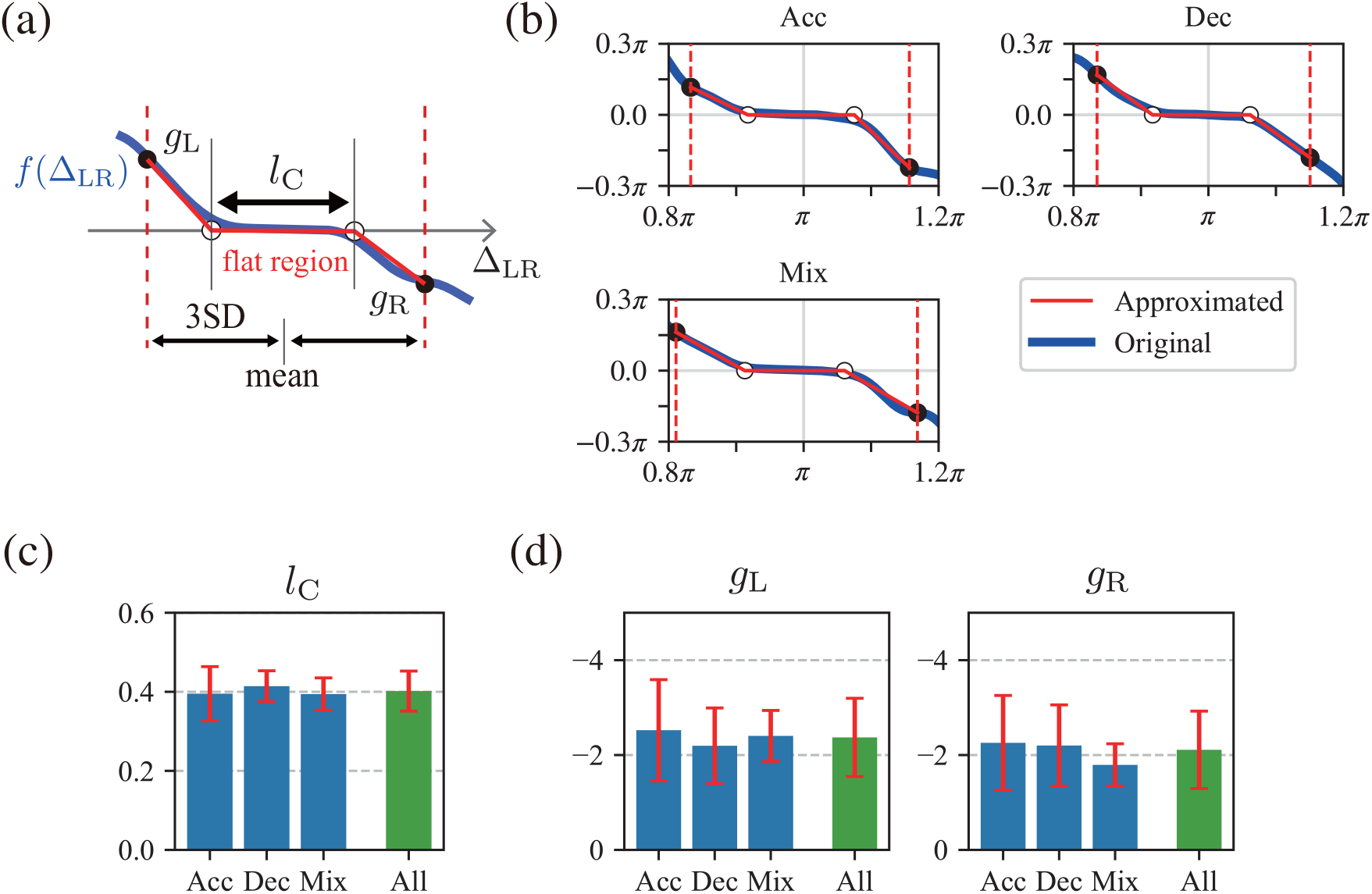
Characteristics of the control of interleg coordination, represented by *f*(Δ_LR_). **(a)** Approximation of *f*(Δ_LR_) by a piecewise linear function within 3 standard deviations (3SD) (vertical dotted lines) from the mean of the observed *f*Δ_LR_. The quantity *l*_C_ is the length of the flat region in which we have *f*(Δ_LR_) = 0, and *g*_L_ and *g*_R_ are the slopes in the left and right outside regions, respectively. The three parameters *l*_C_, *g*_L_, and *g*_R_ were determined by the left and right endpoints of the flat region (white circles) such that the discrepancy between the original and approximated functions is minimized within the region between the two vertical dotted lines under the condition that the approximated function and the original function coincide at the left and right endpoints of the approximated function (black circles). **(b)** Result for the approximation of *f*(Δ_LR_) in Fig. 3(b) under acceleration (Acc), deceleration (Dec), and mixed (Mix) conditions. **(c)** Mean and standard deviation of *l*_C_ among the subjects, where “All” indicates the statistical result from all perturbation conditions. **(d)**Mean and standard deviation of *g*_L_ and *g*_R_ among the subjects.

## Discussion

Using time-series data measured during walking under perturbations that disturbed the antiphase relationship between the motion of the left and right legs (Fig. 2), we constructed a function that models the control of interleg coordination in human walking on the basis of the phase reduction theory and Bayesian inference method. We found that this function has a flat region with values close to 0 in the vicinity of the antiphase condition and steep negative slopes outside this flat region for all subjects studied (Fig. 4). This result indicates that the relative phase between the motion of the left and right legs is not actively controlled until the deviation from the antiphase condition exceeds a certain threshold. In other words, there is a dead zone like that in the case of the steering wheel of an automobile. This differs from our expectation that the relative phase is strictly controlled to maintain the antiphase condition during walking (Fig. 1(c)). This unexpected finding was obtained for the first time through the quantitative evaluation of the control of interleg coordination as a function of the relative phase.

It is plausible that such a forgoing of control reduces the energy consumption during human walking. In addition, it enhances maneuverability to change the walking behavior. For example, because walking along a curved path requires left-right asymmetry in the walking behavior and a phase shift from the antiphase condition [7], the lack of control allows a change in the walking direction. However, large deviations from the antiphase condition result in a deterioration of gait performance. For this reason, when the deviation exceeds a certain threshold, control is activated to reduce this deviation. Although the slopes of *f*(Δ_LR_) outside the flat region vary among the subjects (Fig. 4(d)), the length of the flat region, which determines the threshold beyond which the control is exercised, does not (Fig. 4(c)). This suggests that the threshold is a universal characteristic inherent in the normal walking of healthy young people.

The present result was obtained by determining the specific shape of the function controlling the interleg coordination. The method employed here relies to a large extent on recent developments in the identification of the phase equation using big data [20]. In previous studies [8, 28], interleg coordination has also been investigated through use of the phase equation. However, in those works, a simple form (for example a sine function) was assumed for the coupling function in order to determine the function controlling the interleg coordination. In that case, there appeared no region in which control is not exercised.

The forgoing of control in specific situations in which we expect strict control also appears during quiet standing in humans. Because the upright standing posture is inherently unstable, like an inverted pendulum, it is natural to conjecture that posture is strictly controlled, in order to maintain quiet standing, as we also conjectured that the relative phase of leg motion is strictly controlled in order to maintain the antiphase condition during walking. However, it has been suggested that posture is not actively controlled until deviation from the quiet, upright state exceeds a state-dependent threshold [3, 5, 18, 27]. It thus appears that interleg coordination during walking and quiet standing have a common motor control strategy in humans.

Although the present study focused only on the normal walking of healthy young people, our method has great potential for broader application. For example, because our method can determine the phase coupling functions Γ_LR_ and Γ_RL_ independently, it is applicable to the investigation of the control of interleg coordination during walking with left-right asymmetry, for example in the case of a curved path [7] or a split-belt treadmill [26], or in the case of hemiplegic stroke patients [30]. In addition, the relative phase between the motion of the left and right legs during walking in elderly people and patients with Parkinson’s disease fluctuates significantly [23, 24, 25, 29]. It has been suggested that, unlike young, healthy people, elderly people and patients with Parkinson’s disease do exercise strict control in quiet standing [28]. We plan to investigate how the forgoing of control in interleg coordination during walking changes in elderly people and patients with Parkinson’s disease. We believe that our method will contribute to the understanding of not only motor control strategies employed by young, healthy humans, but also motor disorder mechanisms in elderly people and patients with neurological disabilities. Finally, we point out that our method derives results using data measured when the relative phase between the legs recovers after being disturbed by intermittent perturbations of short duration. At present, this approach has the limitation that it does not allow us to increase the number of perturbations within a period and force subjects to walk for a long time. In the near future, we will improve our method to reduce the burden on subjects and develop applications for the diagnosis and treatment of gait irregularities.

## Methods

### Measurement

To extract the phase coupling functions that govern the control of interleg coordination as modeled by the phase equation, we use kinematic data measured during walking of eight subjects. However, we frequently encountered difficulty in extracting these functions in the case of steady-state walking under normal conditions, because in such situations, the left and right legs are highly synchronized in the antiphase relationship, and for this reason, the measured data provide little information regarding the interaction between the legs. To overcome this difficulty, we varied the belt speed of the treadmill on which the subjects walked in order to disrupt the antiphase relationship between the legs. In response to such perturbations, the control of interleg coordination was exercised, and this allowed us to extract the phase coupling functions from the measured data. This study was approved by the Ethics Committee of Doshisha University. Written informed consent was obtained from all subjects after the procedures had been fully explained. A part of the measured data was used in Refs. [10, 11] for purposes different from that of the present study.

The subjects walked on a treadmill (ITR3017, BERTEC corporation) with a belt speed of 1.0 m/s. We used a motion capture system (MAC3D Digital RealTime System; NAC Image Technology, Inc.) to measure the kinematic data with a sampling frequency of 500 Hz. Reflective markers were attached to the subjects’ skin at several easily identifiable positions on both the left and right sides: the head, the top of the acromion, the greater trochanter, the lateral condyle of the knee, the lateral malleolus, the head of the second metatarsal, and the heel. We measured the belt speed of the treadmill using a rotary encoder with a sampling frequency of 500 Hz.

We applied short duration intermittent perturbations to the walking behavior by suddenly changing the belt speed. We used two types of perturbations: acceleration and deceleration. The acceleration and deceleration perturbations increased and decreased the belt speed, respectively, by 0.6 m/s over a period of 0.1 s. The belt speed then returned to the original speed of 1.0 m/s in the following 0.1 s. The perturbations began approximately 10 s after the start of the measurement process and were applied approximately every 5 s thereafter. Each trial included ten perturbations and lasted approximately 60 s. We used three types of perturbation conditions for each trial: acceleration, deceleration, and mixed conditions. Under the first two conditions, only acceleration and deceleration perturbations, respectively, were applied, while under the mixed condition, both types of perturbations were applied randomly.

All subjects (Subjects A–H) were healthy men (*n* = 8; age 21–23 years; weight 50.2–79.5 kg; height 161–182 cm). Subjects A and B performed 25 trials under both acceleration and deceleration conditions (with the exception that Subject A performed 24 trials under the deceleration condition). The other six subjects (Subjects C–H) performed 15 trials under each of the perturbation conditions. Each trial consisted of between 805 and 1257 walking cycles in the case of acceleration conditions, between 781 and 1248 walking cycles in the case of deceleration conditions, and between 787 and 852 walking cycles in the case of mixed conditions.

### Data processing

In preparation for fitting the various quantities appearing in the phase equation with the measured time-series data, we first transformed the time series of the kinematic angle *φ_i_*(*t*) to those of the oscillator phase, *ϕ_i_*(*t*). (We use the subscript *i* ∈ (L, R), which indicates the left or right leg, instead of L and R in the subsequent sections.) In addition, we transformed the time series of the external perturbation *p*(*t*) to those for the rescaled external perturbation *I*(*t*).

#### Oscillator phase

First, we obtained the time series for the protophase *θ_i_*(*t*) from the time series for the kinematic angle *φ_i_*(*t*) using the Hilbert transform [22] of *φ_i_*(*t*), 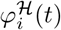, as

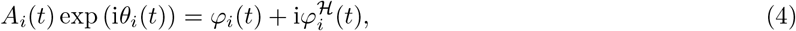

where *A_i_*(*t*) is the amplitude. The phase of an uncoupled limit-cycle oscillator is generally defined in phase reduction theory as always increasing at a constant natural frequency (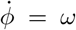 for phase *ϕ* and natural frequency *ω*) [9, 14, 17, 31]. However, the protophase *θ_i_*(*t*) in Eq. (4) does not exhibit such a linear time evolution and is not suitable for the phase equation. To statistically rectify this problem, we transformed *θ_i_*(*t*) to *ϕ_i_*(*t*) as described in Refs. [15, 16] such that it tends to evolve linearly in time with the natural frequency through the definition

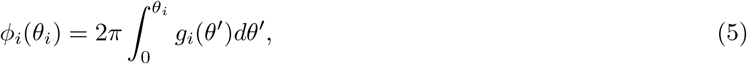

where *g_i_*(*θ_i_*) is the probability density function of *θ_i_*(*t*) calculated from the time-series data for all trials of the walking experiments for each subject and each perturbation condition. We used *ϕ_i_*(*t*) as the oscillator phase.

#### External perturbation

To make the phase sensitivity function *Z_i_* a dimensionless quantity, we introduced *I*(*t*) through the definition 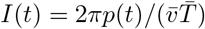, where 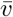 is the average treadmill belt speed, 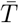 is the average gait cycle duration, and 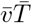 is the average stride length for one gait cycle.

#### Outlier exclusion

Unexpected events during walking, such as stumbling, yields outliers in the derivative of the time series of the oscillator phase. This reduces the accuracy of the fitting. To address this problem, we evaluated the derivative of the oscillator phase at time *t_τ_*, 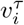, as follows:

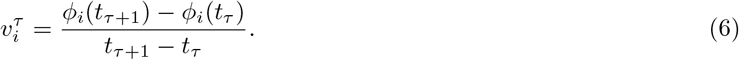

We performed the Smirnov-Grubbs test on the sets of 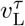 and 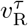 to determine the times at which outliers occur (*p* < 0.05) from the time-series data of all trials. In this procedure, we excluded time series during which external perturbations were applied (green regions in Fig 2) from consideration. If an outlier appeared at some time *t_τ_*, the data for *ϕ_i_*(*t_τ_*) and *I*(*t_τ_*) were excluded from the original data set. Outliers were often detected during touch-down events, when the belt speed exhibited small, sudden variations (Fig 2).

### Fitting the phase equations

Using the time series of the oscillator phase *ϕ_i_*(*t*) and external perturbation *I*(*t*), we fit the parameters in the phase equations (Eqs. (1) and (2)) for each subject and perturbation condition. (We obtained 22 sets of results, corresponding to two perturbation conditions for two subjects and three perturbation conditions for six subjects.) Specifically, we obtained values for the natural frequency *ω_i_*, the phase coupling function Γ*_ij_*, and the phase sensitivity function *Z_i_* on the basis of the Bayesian inference method [17, 19, 20, 21].

#### Phase coupling function

Because the phase coupling function Γ_*ij*_(*ψ*) is 2*π*-periodic, it can be represented by a Fourier series as

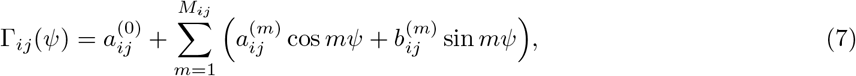

where 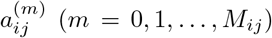 and 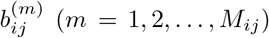 are coefficients that determine the shape of the phase coupling function, and *M_ij_* is the maximum order of the Fourier series. When *M_ij_* = 0, only the constant term remains, and we have 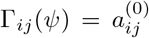. An appropriate value is necessary for *M_ij_*, and it is determined through a model selection, as explained below. To avoid redundancy of the two constant terms *ω_i_* and 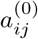 in Eqs. (1), (2), and (7), we define 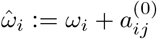 and 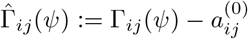.

#### Phase sensitivity function

Because the phase sensitivity function *Z_i_*(*ψ*) is also 2*π*-periodic, it can be represented by a Fourier series as

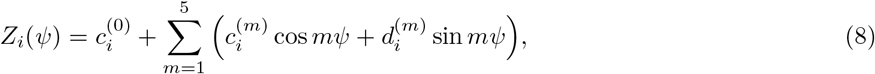

where 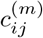 (*m* = 0, 1, …, 5) and 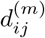 (*m* = 1, 2, …, 5) are coefficients that determine the shape of the phase sensitivity function. Because the phase sensitivity function is often composed of lower-order terms, we used a maximum order of 5. We confirmed that the phase sensitivity functions obtained with this restriction were almost identical to those obtained using higher maximum orders up to 10.

#### Bayesian inference method

Substituting Eqs. (7) and (8) into the phase equations (1) and (2), we obtain

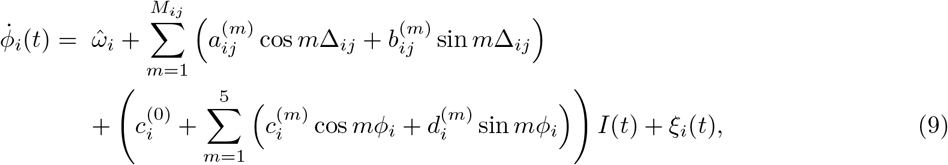

where Δ_*ij*_ = *ϕ_j_* – *ϕ_i_* ((*i, j*) = (L, R), (R, L)). Then, using the time series of the oscillator phase {*φ_i_*(*t_τ_*)}_*i,τ*_ and those of the external perturbation {*I*(*t_τ_*)}_*τ*_ (*τ* = 0, 1,…, *T*), where *T* is the sample number for each subject and perturbation condition, we obtain the unknown parameters 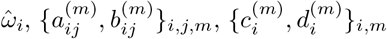, and 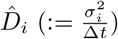, where Δ*t* is the sampling interval. For simplicity, we define the following:

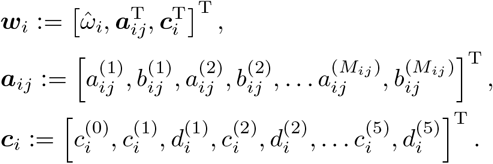

Here, ***ω**_i_*, whose dimension is *N_i_* (≔ 12 + 2*M_ij_*), represents all of the unknown parameters of one phase equation, except for 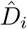, and ***a**_ij_* and ***c**_i_* represent the unknown parameters of the phase coupling function, 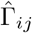, and phase sensitivity function, *Z_i_*, respectively.

Following the Bayesian estimation method [4, 20], we define the likelihood function using a Gaussian distribution 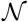 as follows:

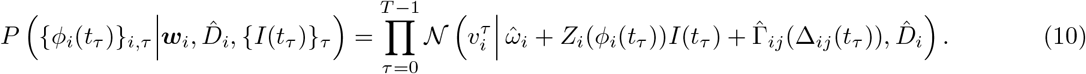

This function represents the probability to reproduce the time series {*ϕ_i_*(*t_τ_*)}_*i,τ*_ when the parameters ***w**_i_* and 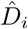 and the time series {*I*(*t_τ_*)}_*τ*_ are given. We adopt a Gaussian-inverse-gamma distribution 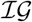 for the conjugate prior distribution as follows:

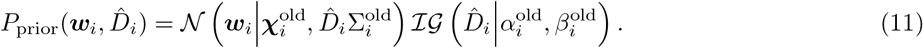

Here, the hyperparameters ***χ***^old^ and Σ^old^ determine the mean and covariance matrices, respectively, of the Gaussian distribution 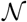, and the other hyperparameters, *α*^old^ and *β*^old^, determine the shape and scale, respectively, of the Gaussian-inverse-gamma distribution 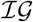, which is given by

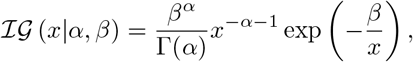

where Γ(·) is the gamma function. In particular, the first element of 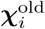 controls 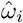, and the other elements control ***a**_ij_* and ***c**_i_*. In this study, the hyperparameters in the prior distribution were set

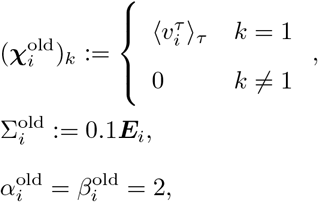

as where ***E**_i_* is the *N_i_* × *N_i_* identity matrix. According to Bayes’ theorem, we obtain the posterior distribution of ***ω**_i_* and 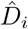 from the product of the likelihood function and prior distribution:

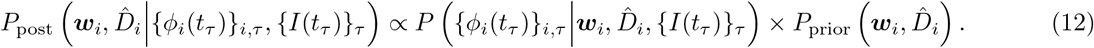

#### Update rule for hyperparameters

Because of the conjugacy of the prior distribution (Eq. (11)) to the likelihood function (Eq. (10)), the posterior distribution (Eq. (12)) is obtained as a Gaussian-inverse-gamma distribution 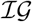 as follows:

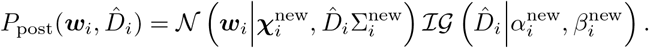

Therefore, it is characterized by the hyperparameters 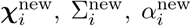, and 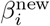 in the same way as the prior distribution (Eq. (11)).

The update rule for the hyperparameters is derived in a simple way in accordance with the Bayesian update rules. First, we define the matrix 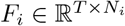 and the vector 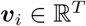 as follows:

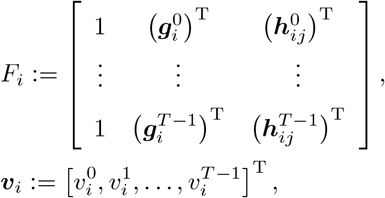

where the vectors 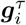 and 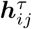 are given by

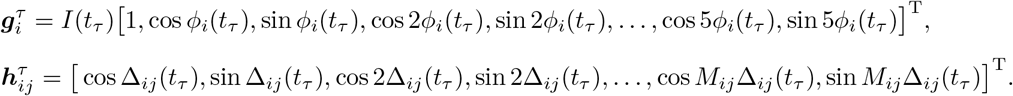

Then, the hyperparameters in the posterior distribution are calculated as follows:

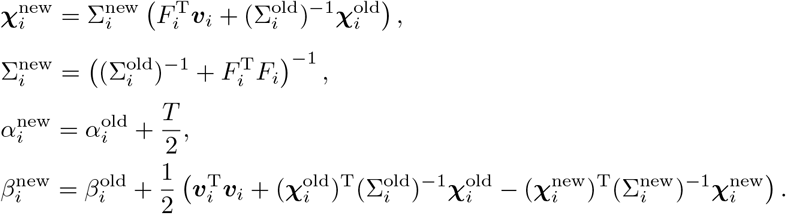

Finally, the unknown parameters ***ω**_i_* and 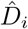 of the phase equation (Eq. (9)) are determined from the mean of the posterior distribution.

#### Model selection

For the phase coupling function (Eq. (7)), we need to choose the model complexity, i.e., an optimal maximum order *M_ij_* of the Fourier series. For example, if *M_ij_* is too small, there will not be sufficiently many terms to correctly represent the function, whereas if *M_ij_* is too large, there will unnecessarily be higher-order harmonic terms involved in representing the function, and this will lead to overfitting. Following standard Bayesian methods [4], we determined an optimal value for *M_ij_* under the assumption that this value maximizes the model evidence *P*_ME_ for 0 ≤ *M_ij_* ≤ 20 as follows:

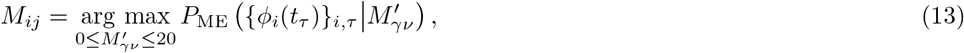

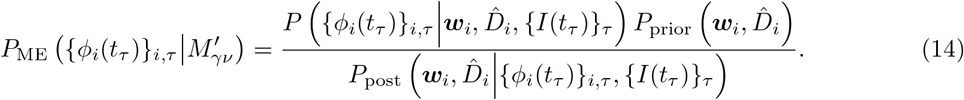

The values of Δ_LR_ are distributed within a narrow region near *π*, and the description provided by the phase coupling function is limited to this region. The lack of data outside this region results in the overfitting. Using this method, the optimal value for *M_ij_* was determined in each case to be in the range 11-20.

## Acknowledgement

This work was supported in part by the following: JSPS KAKENHI Grant Numbers JP20K21810, JP20H04144, and JP20K20520; JST CREST Grant Number JPMJCR09U2; and JST FOREST Program Grant Number JPMJFR2021.

## Author contributions

T. Aoyagi and S. Aoi developed the study design. T. Funato performed human experiments. T. Arai and K. Ota analyzed the data in consultation with K. Tsuchiya, T. Aoyagi, and S. Aoi. T. Arai and S. Aoi wrote the manuscript, and T. Aoyagi revised it. All authors reviewed and approved the final version of the manuscript.

