## Supplementary information for "Interleg coordination is not strictly controlled during walking"

#### A Fitting results for all subjects

Figure S1 displays the fitting results for $Z_{\mathrm{L}}$, $Z_{\mathrm{R}}$, and $f(\Delta_{\mathrm{LR}})$ for all subjects (Subjects A–H) and all perturbation conditions (acceleration, deceleration, and mixed conditions). For all subjects and conditions, both $Z_{\mathrm{L}}$ and $Z_{\mathrm{R}}$ possess a unimodal shape with a peak near $\pi$ and are close to zero in the range from 0 to $\pi/2$. The function $f(\Delta_{\mathrm{LR}})$ has a flat region with a value of 0 near $\pi$ and steep negative slopes outside this flat region. These results are consistent with the result for the representative subject (Subject G), appearing in Fig. 3.

#### B Comparison of fitting results in cases with and without a phase sensitivity function

We incorporated $Z_i I$ ($i \in (\mathrm{L}, \mathrm{R})$) when we derived the control of interleg coordination, $f(\Delta_{\mathrm{LR}})$, from the phase equations (Eqs. (1) and (2)), in contrast with previous studies [1, 2, 3]. To clarify the contribution of $Z_i I$ to the fitting result, we compared the results for $f(\Delta_{\mathrm{LR}})$ obtained in the cases with and without $Z_i I$. We used the following phase equations for the case without $Z_i I$:

$$\dot{\phi}_{\mathrm{L}}(t) = \omega_{\mathrm{L}} + \Gamma_{\mathrm{LR}}(\phi_{\mathrm{R}} - \phi_{\mathrm{L}}) + \xi_{\mathrm{L}}(t), \tag{S1}$$

$$\dot{\phi}_{\mathrm{R}}(t) = \omega_{\mathrm{R}} + \Gamma_{\mathrm{RL}}(\phi_{\mathrm{L}} - \phi_{\mathrm{R}}) + \xi_{\mathrm{R}}(t), \tag{S2}$$

which are simply identical to Eqs. (1) and (2) with $Z_i I$ subtracted out. Because these equations cannot describe the changes caused by external perturbations, we excluded the time series $\phi_i(t)$ ($i \in (\mathrm{L}, \mathrm{R})$) during external perturbations in the fitting (green regions in Fig 2). Then, because outliers often appeared at the start and end of the time series due to edge artifacts of the Hilbert transformation, we removed them using the Smirnov-Grubbs test ($p < 0.05$).

Figure S2 compares the fitting results for $f(\Delta_{\mathrm{LR}})$ in the cases with and without $Z_i I$ for all subjects (Subjects A–H) and all perturbation conditions (acceleration, deceleration, and mixed conditions). Although in both cases, $f(\Delta_{\mathrm{LR}})$ has a flat region with a value of 0 near $\pi$ and steep negative slopes outside this flat region for all subjects and conditions, the fitting results with $Z_i I$ display these characteristics more sharply than those without $Z_i I$.

### C Approximation results for all subjects

Figure S3 displays the approximation results for $f(\Delta_{\mathrm{LR}})$ using a piecewise linear function within 3 standard deviations from the mean of the observed $\Delta_{\mathrm{LR}}$ for all subjects (Subjects A–H) and all perturbation conditions (acceleration, deceleration, and mixed conditions). For all subjects and conditions, the approximation of $f(\Delta_{\mathrm{LR}})$ possesses a flat region with a value of 0 near $\Delta_{\mathrm{LR}} = \pi$ and steep negative slopes outside this flat region. These results are consistent with the result for the representative subject (Subject G), appearing in Fig. 4.

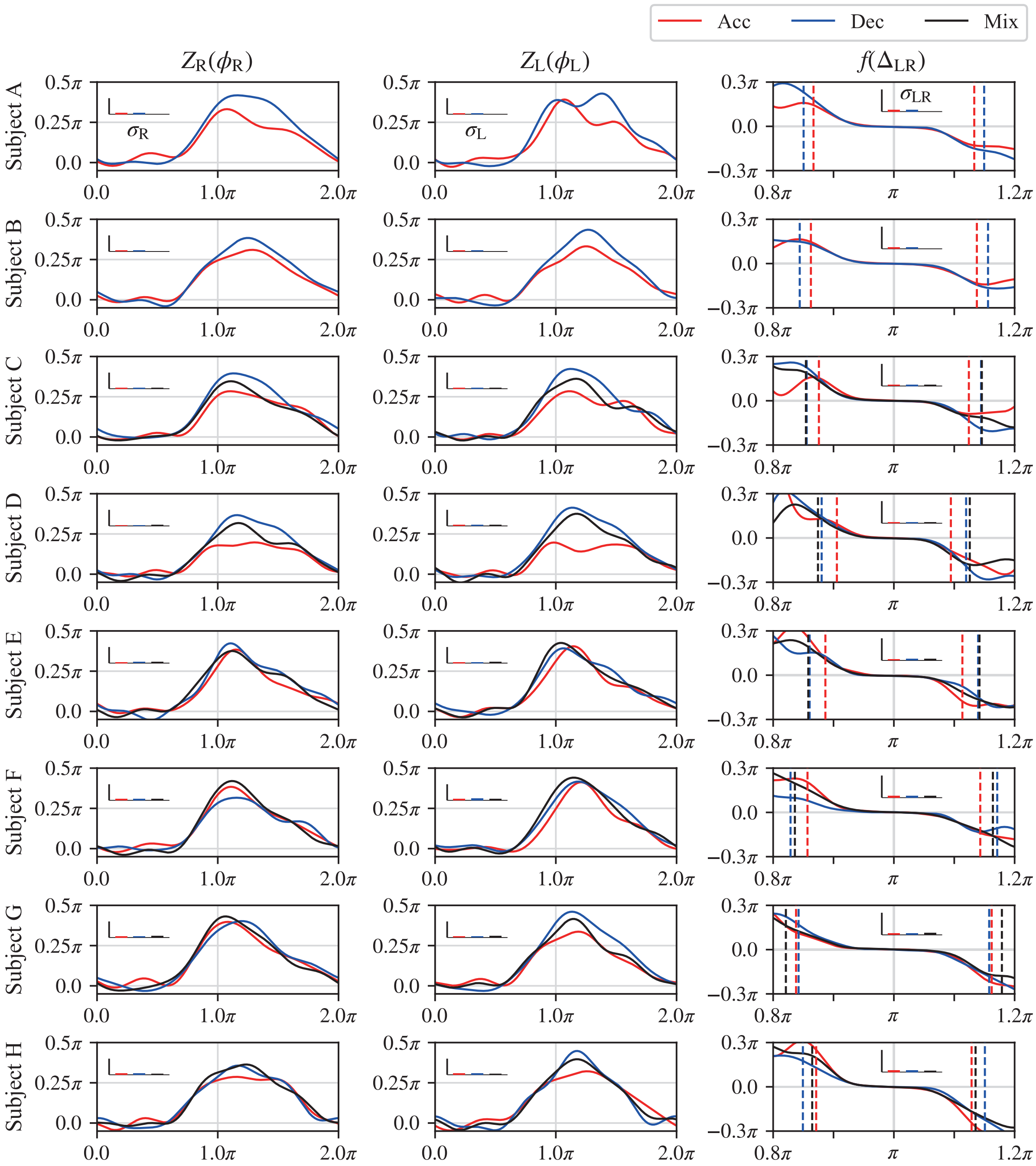


Figure S1: **Fitting results for all subjects.** Phase sensitivity functions $Z_i(\phi_i)$ ($i \in (\mathrm{L}, \mathrm{R})$) and the control of interleg coordination function, $f(\Delta_{\mathrm{LR}})$, in the cases of acceleration (Acc), deceleration (Dec), and mixed (Mix) conditions for all subjects (Subjects A–H). (Note that Subjects A and B performed trials only under the acceleration and deceleration conditions.) The function $f(\Delta_{\mathrm{LR}})$ was shifted to place the mean of the observed values of $\Delta_{\mathrm{LR}}$ at $\pi$ to aid visualization. The vertical dotted lines indicate 3 standard deviations from the mean of the observed values of $\Delta_{\mathrm{LR}}$. The histograms for $Z_{\mathrm{L}}(\phi_{\mathrm{L}}), Z_{\mathrm{R}}(\phi_{\mathrm{R}})$, and $f(\Delta_{\mathrm{LR}})$ display the noise intensities $\sigma_{\mathrm{L}}, \sigma_{\mathrm{R}}$, and $\sigma_{\mathrm{LR}}$, respectively. (The height of each histogram is too small to see.) The mean and standard deviation of the derived noise intensities for all results are $0.0106\pi$ and $0.0021\pi$, respectively, which were too small to cause $\Delta_{\mathrm{LR}}$ to deviate from the flat region of $f(\Delta_{\mathrm{LR}})$.

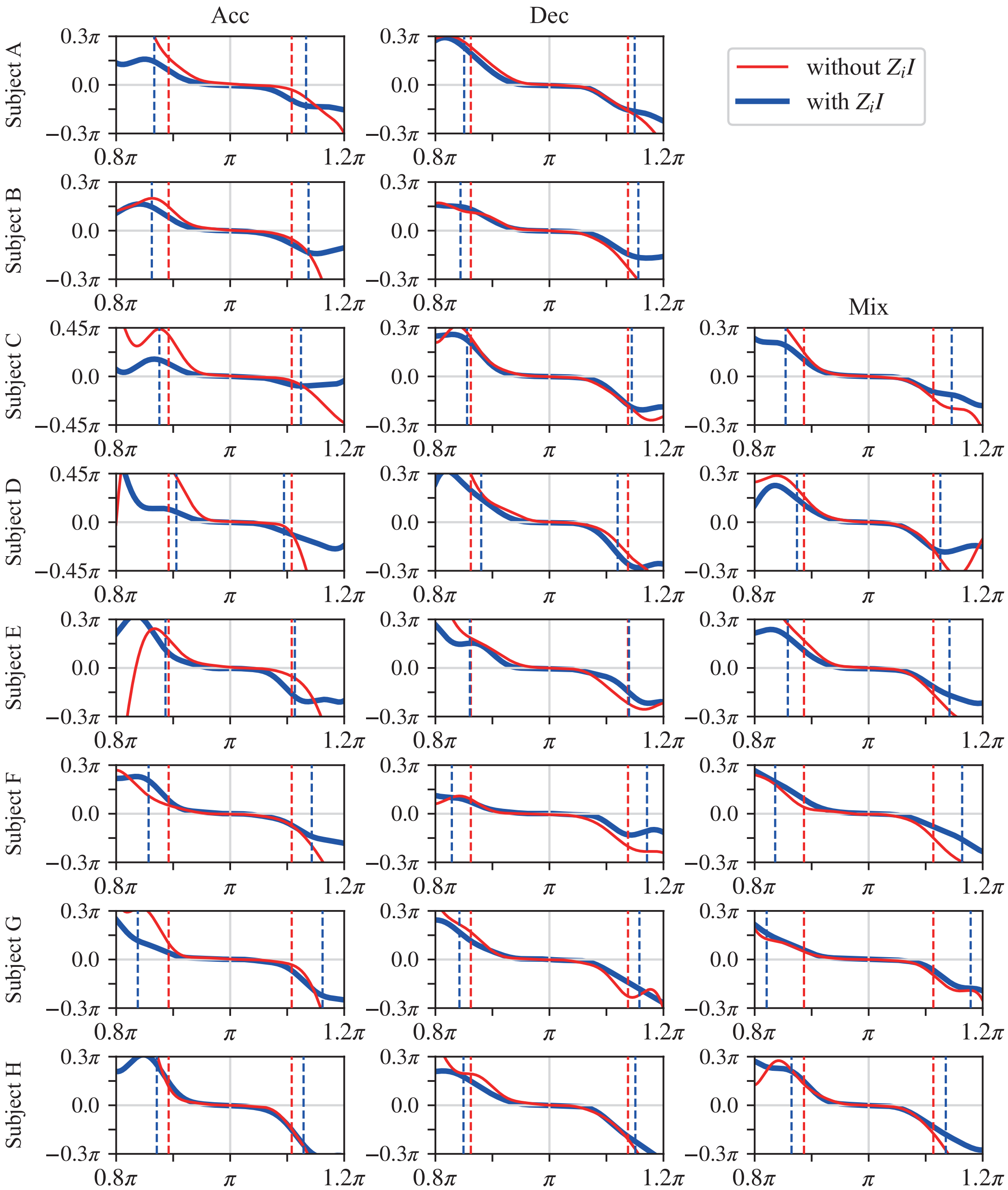


Figure S2: **Comparison of fittings in cases with and without a phase sensitivity function.** $f(\Delta_{\mathrm{LR}})$ in the cases of acceleration (Acc), deceleration (Dec), and mixed (Mix) conditions for all subjects (Subjects A–H). (Note that Subjects A and B performed trials only under the acceleration and deceleration conditions.) $f(\Delta_{\mathrm{LR}})$ was shifted to place the mean of the observed values of $\Delta_{\mathrm{LR}}$ at $\pi$ to aid visualization. The vertical dotted lines indicate 3 standard deviations from the mean of the observed $\Delta_{\mathrm{LR}}$.

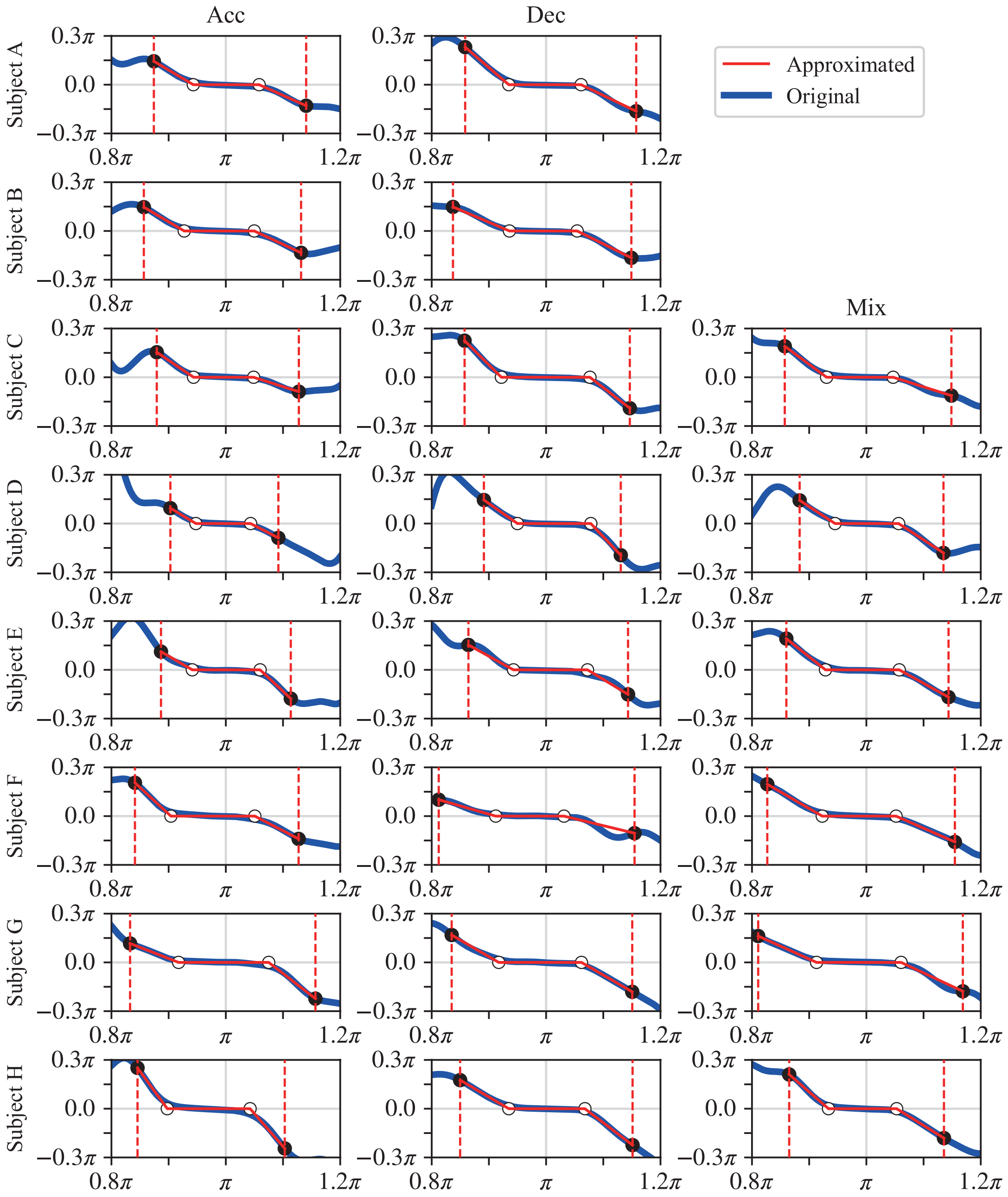


Figure S3: **Approximation results for all subjects.** Approximation of $f(\Delta_{\mathrm{LR}})$ using a piecewise linear function within 3 standard deviations (vertical dotted lines) from the mean of the observed values of $\Delta_{\mathrm{LR}}$ under acceleration (Acc), deceleration (Dec), and mixed (Mix) conditions for all subjects (Subjects A–H). (Subjects A and B performed trials only under the acceleration and deceleration conditions.) The approximated functions were determined by the left and right endpoints of the flat region (white circles) such that the discrepancy between the original and approximated functions was minimized within the region between the two vertical dotted lines under the condition that the approximated function and the original function coincide at the left and right endpoints of the approximated functions (black circles).
